# TOR1 dysfunction promotes phenotypic diversity in the asexual fungus *Candida albicans*

**DOI:** 10.1101/2023.05.12.539650

**Authors:** Lucia F. Zacchi, Jill Rohling, Noah Johnson, Mark T. Anderson, Peter J. Southern, Dana A. Davis

## Abstract

Genetic variation is a primary contributor to phenotypic variation within a population. In asexual eukaryotes however, it is unclear how, or if, genetic variation is generated and maintained to promote phenotypic variation. *C. albicans*, an asexual fungus that causes opportunistic infections in susceptible hosts, has several phenotypic switching systems, including the colony morphology phenotypic switching (CMPS) system. CMPS, a penetrant change in colony morphology on solid medium *in vitro*, is associated with incipient or fulminant clinical disease. CMPS results in the alteration of additional virulence properties, including drug resistance, that are not tightly correlated with colony morphology. Importantly, it is unknown whether CMPS is a regulated or stochastic process. We found that specific mutants affecting the <u>T</u>arget <u>o</u>f <u>R</u>apamycin (TOR) pathway showed an increase in CMPS frequency and CMPS in these mutant backgrounds was associated with changes in rapamycin sensitivity. We also identified growth conditions that promoted CMPS in clinical strains and found that CMPS in these backgrounds was also linked to the TOR pathway through changes in rapamycin sensitivity. These results demonstrate that CMPS promotes phenotypic variation through the TOR pathway, supporting a model that this is a regulated process. Since the TOR growth control pathway is conserved throughout the eukarya, the identification of TOR as a phenotypic diversity regulator likely has broad implications.

## Introduction

Phenotypic diversity within a population can support rapid adaption to new environmental conditions [1]. Phenotypic diversity, an attribute of genetic diversity, is maintained by sexual reproduction in the eukarya. However, numerous unicellular eukaryotes are asexual or predominantly asexual yet generate distinct sub-populations of phenotypic variants. Furthermore, phenotypically distinct sub-populations of cells within tumors have also been described and these sub-populations can impact chemotherapeutic treatment [2]. Recent work also indicates phenotypically distinct sub-populations exist in ostensibly normal tissues [3]. Thus, a key biological question exists, how is phenotypic diversity generated and maintained in asexual populations?

*Candida albicans* is a commensal of human mucosal surfaces and is the most common agent of mucosal and systemic fungal infections [4]. While *C. albicans* is a diploid organism, it lacks a sexual cycle like many fungal pathogens [5][6]. A parasexual cycle has been described, where two diploid cells fuse to form a tetraploid cell, which subsequently reverts back to the diploid state by whole chromosome loss [7]. Tetraploidy, due to the parasexual cycle, can greatly increase the loss of heterozygosity and the formation of aneuploidies leading to genetic diversity [8]. However, population studies indicate that the parasexual cycle occurs rarely in nature [9]. Furthermore, within the human host, much work suggests that there is limited *C. albicans* genetic diversity with a host being colonized primarily with one genetic background [10]. Despite the limited genetic diversity of the *C. albicans* population within a host, phenotypic variants of *C. albicans*, based on colony morphology, have been readily isolated from mucosal surfaces of patients since at least the 1950s [11–13]. These studies demonstrated that phenotypic diversity occurs commonly within the host but did not determine whether this diversity arose from within the limited genetic diversity of the endogenous populations or from neocolonization by genetically distinct strains.

Several studies showed that phenotypic variants can arise in certain mutant backgrounds and during infection in animal models not normally colonized with *C. albicans* [14–17]. These studies suggest that phenotypic diversity arises from within the clonal population. However, it is unclear whether these studies represent analogous diversity generating processes described in the human host.

The colony morphology phenotypic switching (CMPS) system is a well described system of *C. albicans* phenotypic variation occurring within the human host [18]. CMPS is characterized by the formation of colonies with morphologies distinct from the normal smooth cream colored colony. There are three distinct properties that characterize CMPS. First, the morphological variant is penetrant, cells cultured from a colony with a specific morphology give rise to colonies with that morphology. Second, once CMPS initiates it continues at low frequency (∼10^-2^ - 10^-3^). In other words, cells cultured from a colony with a specific morphology can give rise to rare colonies with a new colony morphology including the wild-type morphology [18]. Third, independent CMPS variants with the same colony morphology can differ in other cellular properties, including antifungal drug sensitivity [11]. Thus, the CMPS system generates and maintains phenotypic diversity within a *C. albicans* population.

Unlike analysis of mutant *C. albicans in vitro* or through animal models, the CMPS system was identified *in vivo* in both oral and vaginal niches [11][18]. In these clinical studies, onset of CMPS was observed in conjunction with an increase in population size. For example, CMPS was observed from oral isolates from HIV+ patients with reduced CD4 T-cell counts, but prior to the onset of AIDS or oral candidiasis [11]. Similarly, CMPS was observed in *C. albicans* isolated from women suffering from vaginal yeast infections but not from *C. albicans* recovered from the vaginal tract of non-symptomatic women [13,19]. These clinical studies suggest that CMPS occurs concurrently with an increase in the *C. albicans* population size and may be a predictor for incipient disease.

While much work has been done describing the CMPS phenomenon and characterizing the specific CMPS isolates, several critical questions remain. For example, is CMPS a stochastic process or a regulated process and what is the proximate cause of the phenotypic changes? Since CMPS is correlated with population size, one possibility is quorum sensing systems are involved [20]. However, CMPS has not been observed in mutants affecting known quorum sensing systems in *C. albicans*. Furthermore, the diverse unlinked phenotypes associated with CMPS are reminiscent of random assortment of characteristics observed in Mendelian sexual crosses, but as noted above, sexual reproduction has not been observed in *C. albicans* nor is there population genetic data to suggest such sexual cycles exist. The development of novel diverse phenotypes is also well established in cancer cell populations [2]. Cancer is caused by uncontrolled cellular growth and requires numerous cellular changes including dysfunction of growth promoting pathways, such as the Target of Rapamycin (TOR) and/or Ras signal transduction pathways.

TOR, an essential serine-threonine kinase, is conserved throughout the eukarya and governs cellular growth primarily in response to nitrogen but can also respond to other environmental cues like hormones in mammals [21,22]. In the presence of activating signals, TOR phosphorylates growth promoting effectors, like Sch9 (S6 kinase in mammals) which promotes ribosome biogenesis and translation initiation [23].

Furthermore, active TOR phosphorylates Tap42 inhibiting starvation responses like autophagy and nitrogen catabolite repression [24,25]. In many cancers, TOR dysfunction leads to cellular growth in the absence of activation signals [26]. Furthermore, these TOR-dysfunction dependent cancers readily evolve new phenotypes, including chemotherapeutic drug resistance [27,28].

Here, we report the identification of the TOR growth control pathway as a key regulator of CMPS in *C. albicans*. We found that mutants affecting the TOR pathway promote CMPS and that CMPS isolates in these mutant backgrounds show altered sensitivity to the TOR inhibitor rapamycin. Furthermore, we establish *in vitro* conditions that allow recovery of CMPS variants in wild-type lab and clinical isolates independent of exogenous mutagens and found that these wild-type CMPS isolates also show altered rapamycin sensitivity. Because the TOR pathway is conserved throughout eukaryotic evolution, this may be a conserved mechanism for cellular populations to be able to rapidly generate phenotypic variants in order to respond to novel environmental stresses.

## Methods & Materials

### Media and growth conditions

*C. albicans* strains were routinely grown at 30°C in YPD broth (2% Bacto-peptone, 1% yeast extract, 2% dextrose). For solid YPD medium, 2% Bacto-agar was added [29]. Solid spider medium was made as described previously (1% mannitol, 1% nutrient broth, 0.2% K_2_HPO_4_, pH 7.2 before autoclaving, 1.35% Bacto-agar) [30]. Uridine was added to all media, unless otherwise indicated, to a final concentration of 80µg/ml. Rapamycin (Sigma-Aldrich) was suspended in ethanol and added to solid YPD and spider medium at the indicated concentrations just prior to pouring. Similarly, ethanol was added to solid YPD and spider medium just prior to pouring.

### Strains and plasmids

All yeast strains used in this study are listed in Table 1. SSY switched strains were isolated from papillae of *mds3*Δ*/*Δ colonies using a sterile toothpick and streaking onto YPD plates. SSY parent strains were isolated from a non-papillated region of the same *mds3*Δ*/*Δ colony.

**Table 1.**
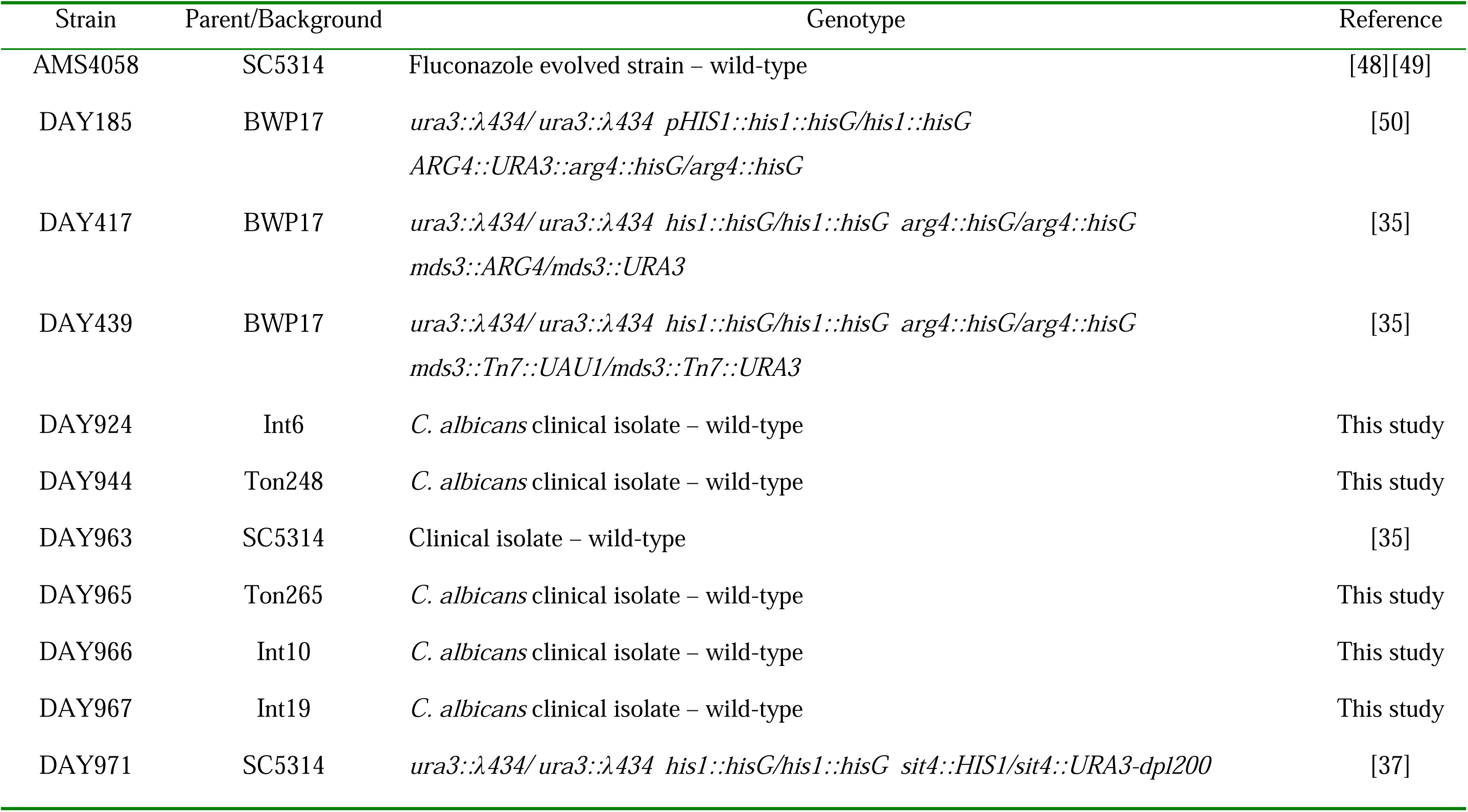
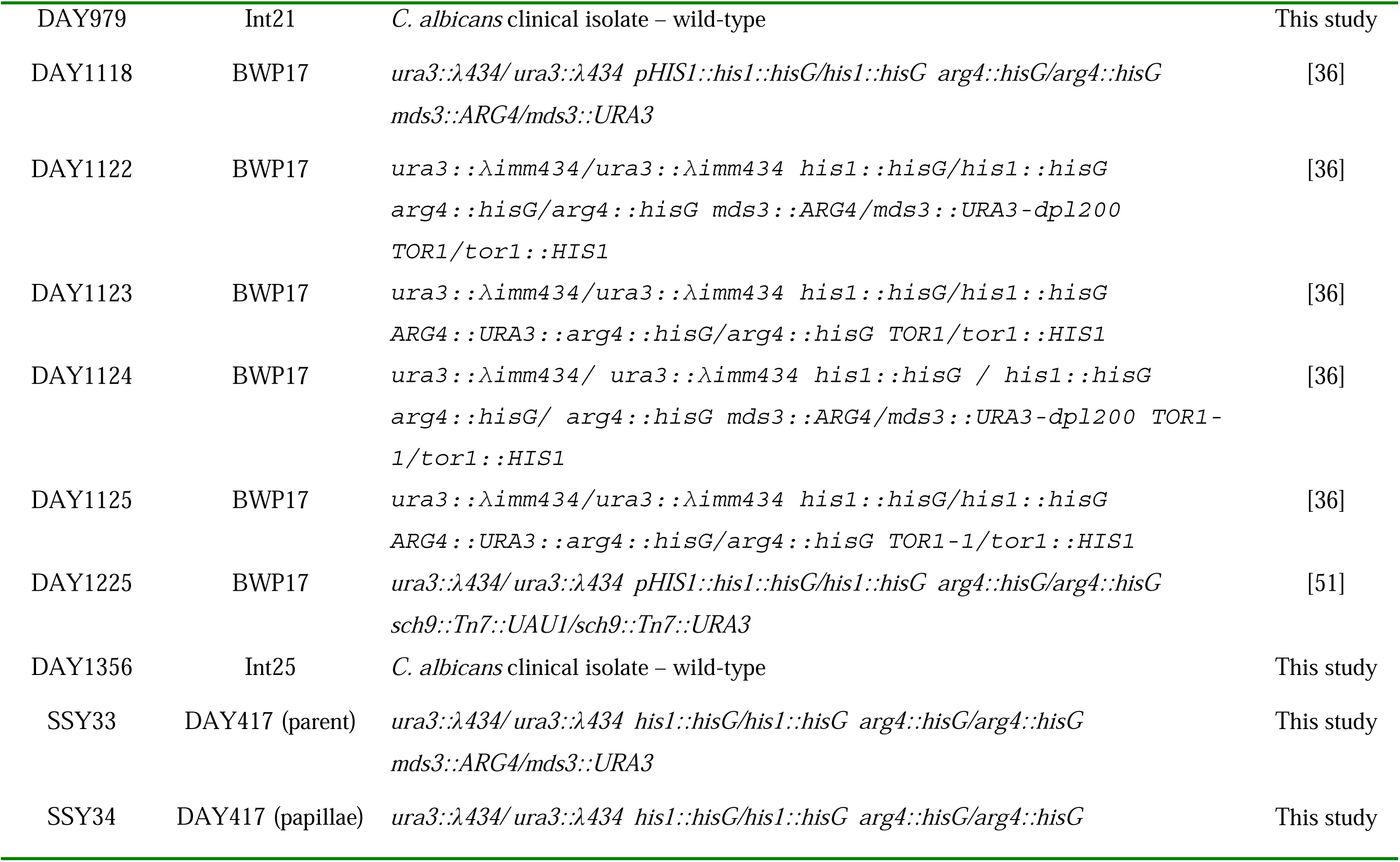
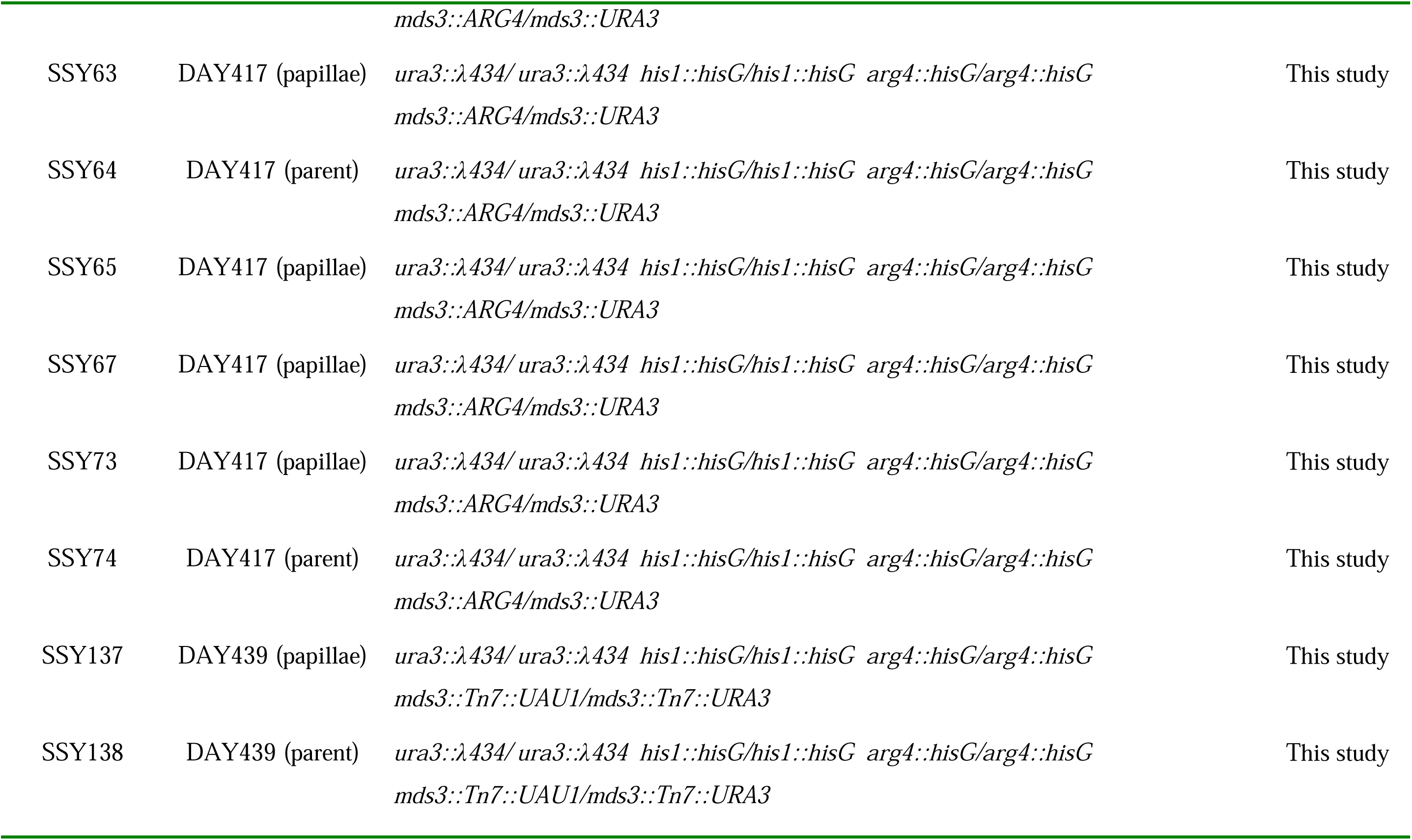
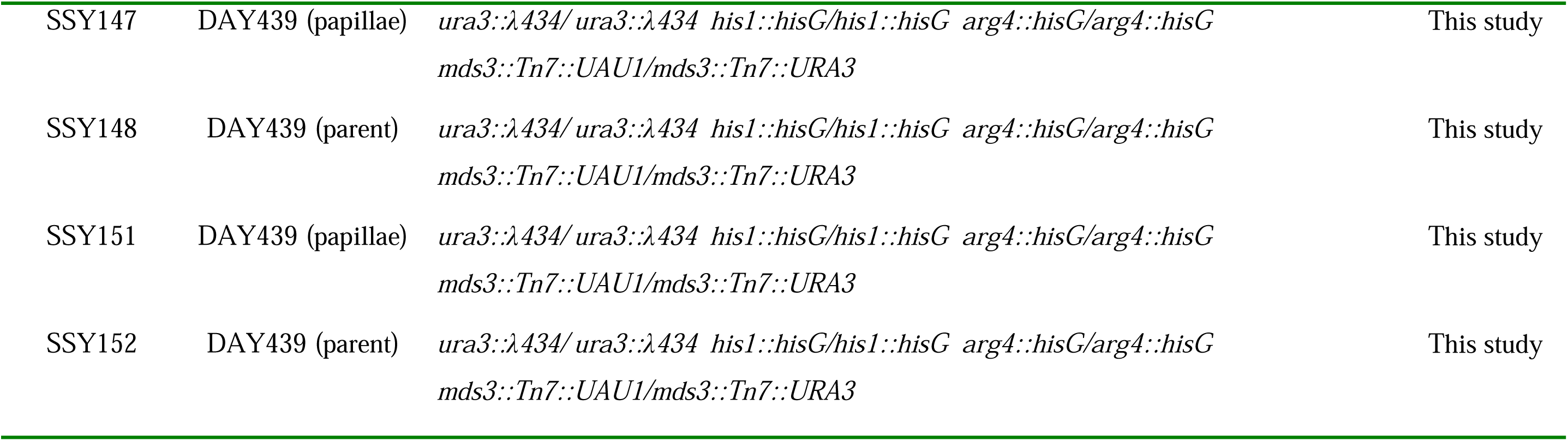
Strains used in this study.

| Strain | Parent/Background | Genotype | Reference |
| --- | --- | --- | --- |
| AMS4058 | SC5314 | Fluconazole evolved strain – wild-type | [48][49] |
| DAY185 | BWP17 | *ura3::λ434/ ura3::λ434 pHIS1::his1::hisG/his1::hisG*<br>*ARG4::URA3::arg4::hisG/arg4::hisG* | [50] |
| DAY417 | BWP17 | *ura3::λ434/ ura3::λ434 his1::hisG/his1::hisG arg4::hisG/arg4::hisG*<br>*mds3::ARG4/mds3::URA3* | [35] |
| DAY439 | BWP17 | *ura3::λ434/ ura3::λ434 his1::hisG/his1::hisG arg4::hisG/arg4::hisG*<br>*mds3::Tn7::UAU1/mds3::Tn7::URA3* | [35] |
| DAY924 | Int6 | *C. albicans* clinical isolate – wild-type | This study |
| DAY944 | Ton248 | *C. albicans* clinical isolate – wild-type | This study |
| DAY963 | SC5314 | Clinical isolate – wild-type | [35] |
| DAY965 | Ton265 | *C. albicans* clinical isolate – wild-type | This study |
| DAY966 | Int10 | *C. albicans* clinical isolate – wild-type | This study |
| DAY967 | Int19 | *C. albicans* clinical isolate – wild-type | This study |
| DAY971 | SC5314 | *ura3::λ434/ ura3::λ434 his1::hisG/his1::hisG sit4::HIS1/sit4::URA3-dpl200* | [37] |
| DAY979 | Int21 | *C. albicans* clinical isolate – wild-type | This study |
| DAY1118 | BWP17 | *ura3::λ434/ura3::λ434 pHIS1::his1::hisG/his1::hisG arg4::hisG/arg4::hisG mds3::ARG4/mds3::URA3* | [36] |
| DAY1122 | BWP17 | *ura3::λimm434/ura3::λimm434 his1::hisG/his1::hisG arg4::hisG/arg4::hisG mds3::ARG4/mds3::URA3-dpl200 TOR1/tor1::HIS1* | [36] |
| DAY1123 | BWP17 | *ura3::λimm434/ura3::λimm434 his1::hisG/his1::hisG ARG4::URA3::arg4::hisG/arg4::hisG TOR1/tor1::HIS1* | [36] |
| DAY1124 | BWP17 | *ura3::λimm434/ura3::λimm434 his1::hisG / his1::hisG arg4::hisG/arg4::hisG mds3::ARG4/mds3::URA3-dpl200 TOR1-1/tor1::HIS1* | [36] |
| DAY1125 | BWP17 | *ura3::λimm434/ura3::λimm434 his1::hisG/his1::hisG ARG4::URA3::arg4::hisG/arg4::hisG TOR1-1/tor1::HIS1* | [36] |
| DAY1225 | BWP17 | *ura3::λ434/ura3::λ434 pHIS1::his1::hisG/his1::hisG arg4::hisG/arg4::hisG sch9::Tn7::UAU1/sch9::Tn7::URA3* | [51] |
| DAY1356 | Int25 | *C. albicans* clinical isolate – wild-type | This study |
| SSY33 | DAY417 (parent) | *ura3::λ434/ura3::λ434 his1::hisG/his1::hisG arg4::hisG/arg4::hisG mds3::ARG4/mds3::URA3* | This study |
| SSY34 | DAY417 (papillae) | *ura3::λ434/ura3::λ434 his1::hisG/his1::hisG arg4::hisG/arg4::hisG* *mds3::ARG4/mds3::URA3* | This study |
| SSY63 | DAY417 (papillae) | *ura3::λ434/ ura3::λ434 his1::hisG/his1::hisG arg4::hisG/arg4::hisG*<br>*mds3::ARG4/mds3::URA3* | This study |
| SSY64 | DAY417 (parent) | *ura3::λ434/ ura3::λ434 his1::hisG/his1::hisG arg4::hisG/arg4::hisG*<br>*mds3::ARG4/mds3::URA3* | This study |
| SSY65 | DAY417 (papillae) | *ura3::λ434/ ura3::λ434 his1::hisG/his1::hisG arg4::hisG/arg4::hisG*<br>*mds3::ARG4/mds3::URA3* | This study |
| SSY67 | DAY417 (papillae) | *ura3::λ434/ ura3::λ434 his1::hisG/his1::hisG arg4::hisG/arg4::hisG*<br>*mds3::ARG4/mds3::URA3* | This study |
| SSY73 | DAY417 (papillae) | *ura3::λ434/ ura3::λ434 his1::hisG/his1::hisG arg4::hisG/arg4::hisG*<br>*mds3::ARG4/mds3::URA3* | This study |
| SSY74 | DAY417 (parent) | *ura3::λ434/ ura3::λ434 his1::hisG/his1::hisG arg4::hisG/arg4::hisG*<br>*mds3::ARG4/mds3::URA3* | This study |
| SSY137 | DAY439 (papillae) | *ura3::λ434/ ura3::λ434 his1::hisG/his1::hisG arg4::hisG/arg4::hisG*<br>*mds3::Tn7::UAU1/mds3::Tn7::URA3* | This study |
| SSY138 | DAY439 (parent) | *ura3::λ434/ ura3::λ434 his1::hisG/his1::hisG arg4::hisG/arg4::hisG*<br>*mds3::Tn7::UAU1/mds3::Tn7::URA3* | This study |
| SSY147 | DAY439 (papillae) | *ura3::λ434/ura3::λ434 his1::hisG/his1::hisG arg4::hisG/arg4::hisG*<br>*mds3::Tn7::UAU1/mds3::Tn7::URA3* | This study |
| SSY148 | DAY439 (parent) | *ura3::λ434/ura3::λ434 his1::hisG/his1::hisG arg4::hisG/arg4::hisG*<br>*mds3::Tn7::UAU1/mds3::Tn7::URA3* | This study |
| SSY151 | DAY439 (papillae) | *ura3::λ434/ura3::λ434 his1::hisG/his1::hisG arg4::hisG/arg4::hisG*<br>*mds3::Tn7::UAU1/mds3::Tn7::URA3* | This study |
| SSY152 | DAY439 (parent) | *ura3::λ434/ura3::λ434 his1::hisG/his1::hisG arg4::hisG/arg4::hisG*<br>*mds3::Tn7::UAU1/mds3::Tn7::URA3* | This study |

Commensal clinical isolates were recovered from ostensibly normal tissue specimen derived from colonic (Int) or tonsillar (Ton) tissue (Table 1). Human tissues were obtained and maintained as reported previously [31]. Foci of fungi spontaneously grew on ∼50% of these tissues, but not in controls for fungal contamination. Fungal isolates were purified on YPD medium containing penicillin and streptomycin and grown at 37°C. *C. albicans* isolates were identified using germ tube medium (REMEL) and presumptive candidates confirmed by restriction digest of purified genomic DNA [32] and amplification and sequencing of PCR products of the *ITS1/2* locus [33].

### Growth on rapamycin

For growth assays in the presence of rapamycin, strains were grown overnight in liquid YPD at 30°C. Cells were then serially diluted 10-fold in ddH_2_O in a 96 well microtiter dish and spotted on YPD or Spider medium supplemented with rapamycin or solvent (ethanol) alone using a 48-pin replicator. Plates were incubated at 30°C for 2 or 3 days prior to photography.

### Fluconazole sensitivity

Overnight cultures were grown in YPD at 30°C. The following day, cells were diluted to an OD600 of 0.1 in ddH2O and 100µl cells were spread onto YPD (containing 1% dextrose) using sterile glass beads. A sterile 7mm Whatmann paper disk was placed on the plate and infused with 15 µg of fluconazole suspended in methanol. Plates were wrapped in parafilm and incubated at 30°C for 48 hours prior to photography. Resistance and tolerance numbers were determine using diskImageR [34].

## Results

### Papillae Formation in *mds3*Δ*/*Δ mutants

*MDS3* encodes a non-essential protein that functions downstream of the <u>T</u>arget <u>o</u>f <u>R</u>apamycin (TOR) kinase [35,36]. *C. albicans* mutants lacking Mds3 form smooth round white colonies when grown on solid medium, similar to wild-type cells [36]. However, with prolonged incubation (> 7 days), we observed papillae formation on the surface of *mds3*Δ*/*Δ colonies, but not wild-type colonies (Figure 1A). Papillae formation was observed in both *mds3* insertion and deletion mutants but were not observed in *mds3*Δ*/*Δ *+MDS3* complemented strains even with longer incubation times (data not shown). We did note some hyphal outgrowth from wild-type colonies with increased incubation time (Figure 1A), a known starvation response. Papillae formation increases over time, with < 5% of *mds3*Δ*/*Δ colonies having at least one papillae on day 5 to all colonies having at least one papillae by day 20. These results suggest that Mds3 is a negative regulator of papillae formation.

**Figure 1:**
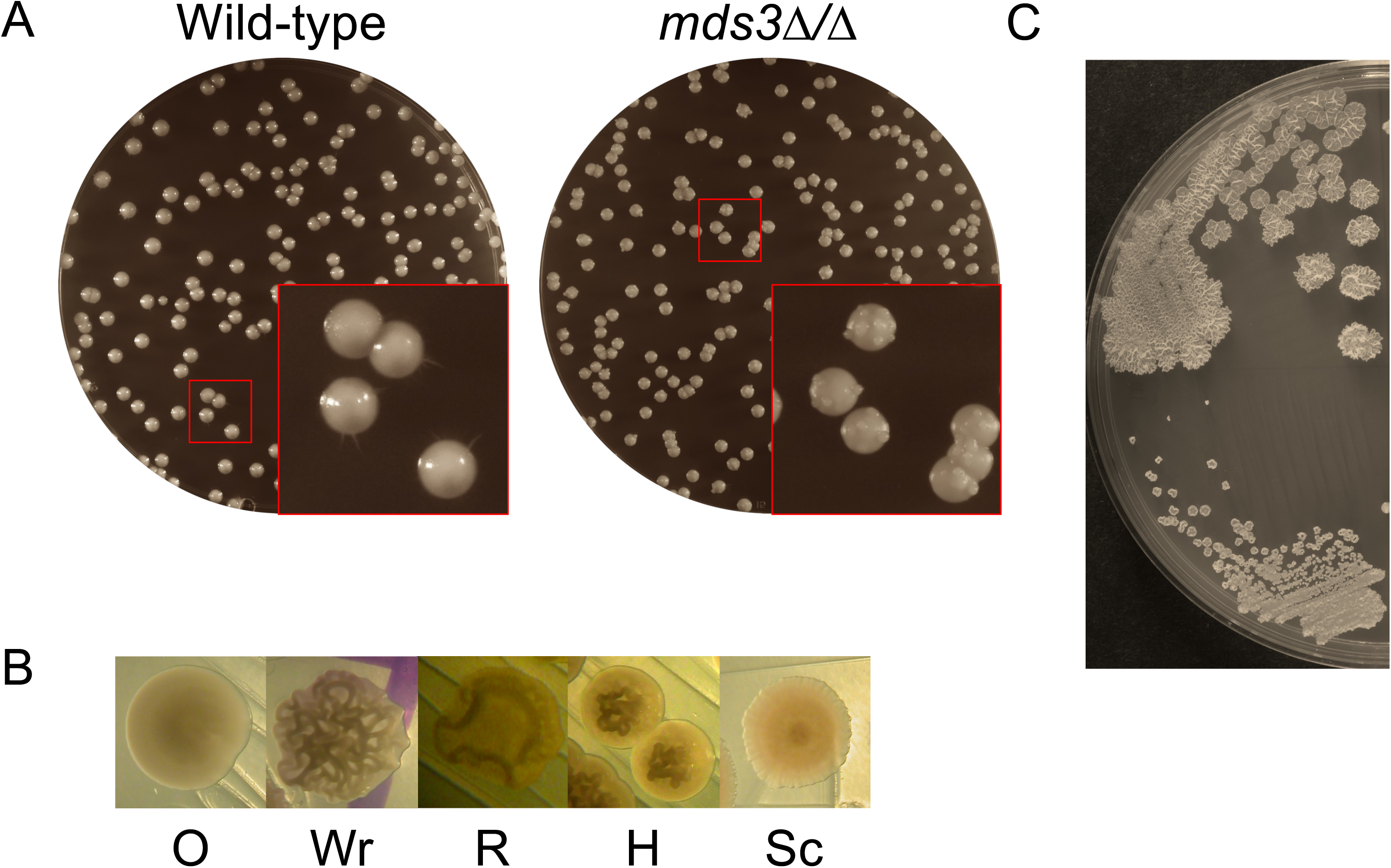
A. Growth of wild-type (DAY185) and a *mds3*Δ*/*Δ mutant (DAY1118) on YPD after prolonged growth. B. Colonies derived from non-papillated regions of DAY1118 give rise to normal smooth cream-colored colonies (O) whereas colonies derived from papillae can give rise to distinct colony morphologies including wrinkled (Wr), ring (R), hat (H), and scalloped (Sc). C. Distinct papillae from the same parent colony can yield distinct colony morphologies. Two distinct papilli were picked from a single colony and streaked for isolation yielding a ‘wrinkled’ and ‘ring’ colony morphology, top and bottom respectively. (B and C) Photographs were taken after 2 days incubation on YPD medium at 37°C.

Because papillae formation represents abnormal growth under starvation conditions not observed in wild-type colonies, we pursued this phenomenon. We first purified cells from papillae to determine if they behaved similarly to the original *mds3*Δ*/*Δ mutant and if not to characterize them. First, purification of cells from the smooth non-papillated portion of the colony resulted in colonies that were phenotypically indistinguishable for the parental *mds3*Δ*/*Δ strain (Figure 1B and 2). This suggests that the age of the colony is not necessarily causing a phenotypic change. Second, colonies derived from papillae generally did not grow like the parental *mds3*Δ*/*Δ strain having differences in colony morphology, colony size, and/or colony color (Figure 1B & Table 2). The most common colony morphology phenotype was the ‘wrinkled’ with the entire surface of the colony covered in ridges (Figure 1B Wr). We also observed colonies with the ‘ring’ phenotype represented by a colony with a single circular ridge near the colony edge, the ‘hat’ phenotype represented by colonies with a series of ridges surrounded by smooth colony, and a ‘scalloped’ phenotype represented by flat colonies with a slightly wavy surface pattern (Figure 1B R, H, and Sc respectively). These colony morphologies are reminiscent of the CMPS variants described previously [18]. Although colonies derived from different papillae can vary from one another, colonies from an individual papilla had the same morphology, size, and color (Figure 1C), demonstrating that the changes occurring to give rise to a papilla are penetrant. Furthermore, restreaking a purified switched colony onto fresh medium gave rise to the same morphology suggesting that the switched phenotype is stable.

**Figure 2:**
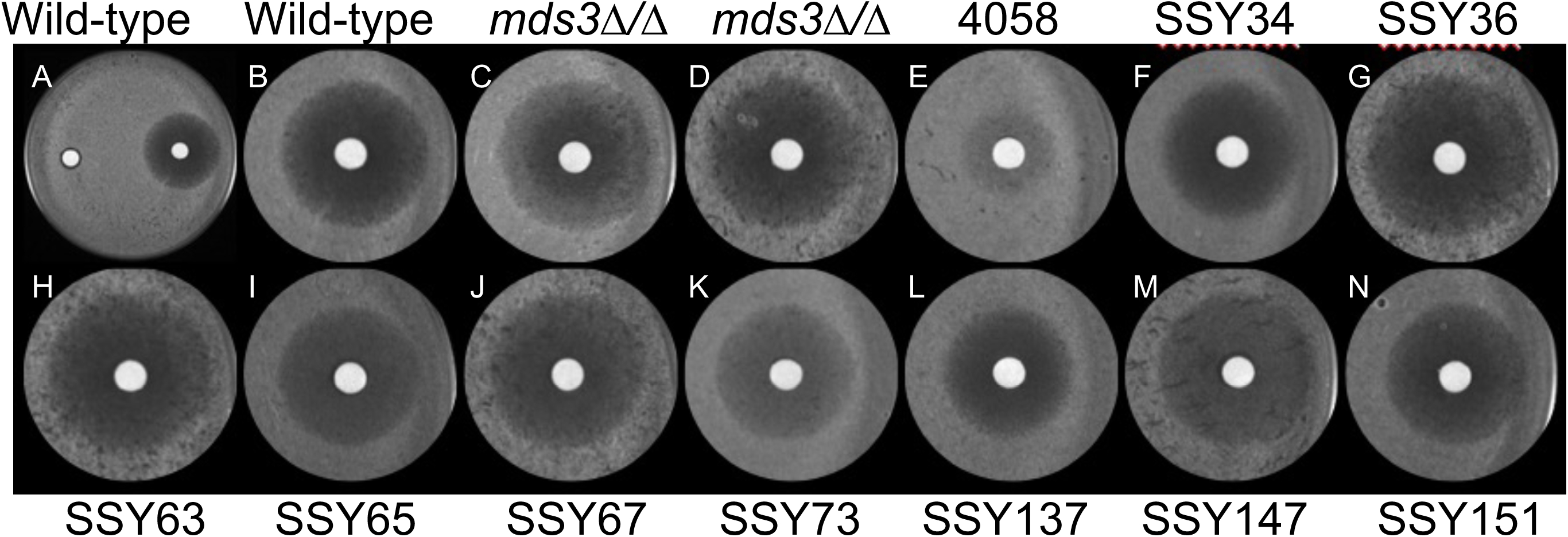
Fluconazole sensitivity and tolerance in independent wrinkled strains. Lawns of each strain were plated on YPD (1% dextrose) and methanol added to a piece of sterile Whatmann paper (left disk) or fluconazole (right disk) and plates were grown at 30°C for 48 hours. (A) Wild-type (DAY286) plate. (B-N) Close up of the fluconazole disk for each strain (B, wild-type close up of A); C and D *mds3*Δ*/*Δ mutants (DAY417 and DAY439 respectively); E AMS4058 (wild-type fluconazole evolved strain); F-N *mds3*Δ*/*Δ derived CMPS strains.

**Table 2:**
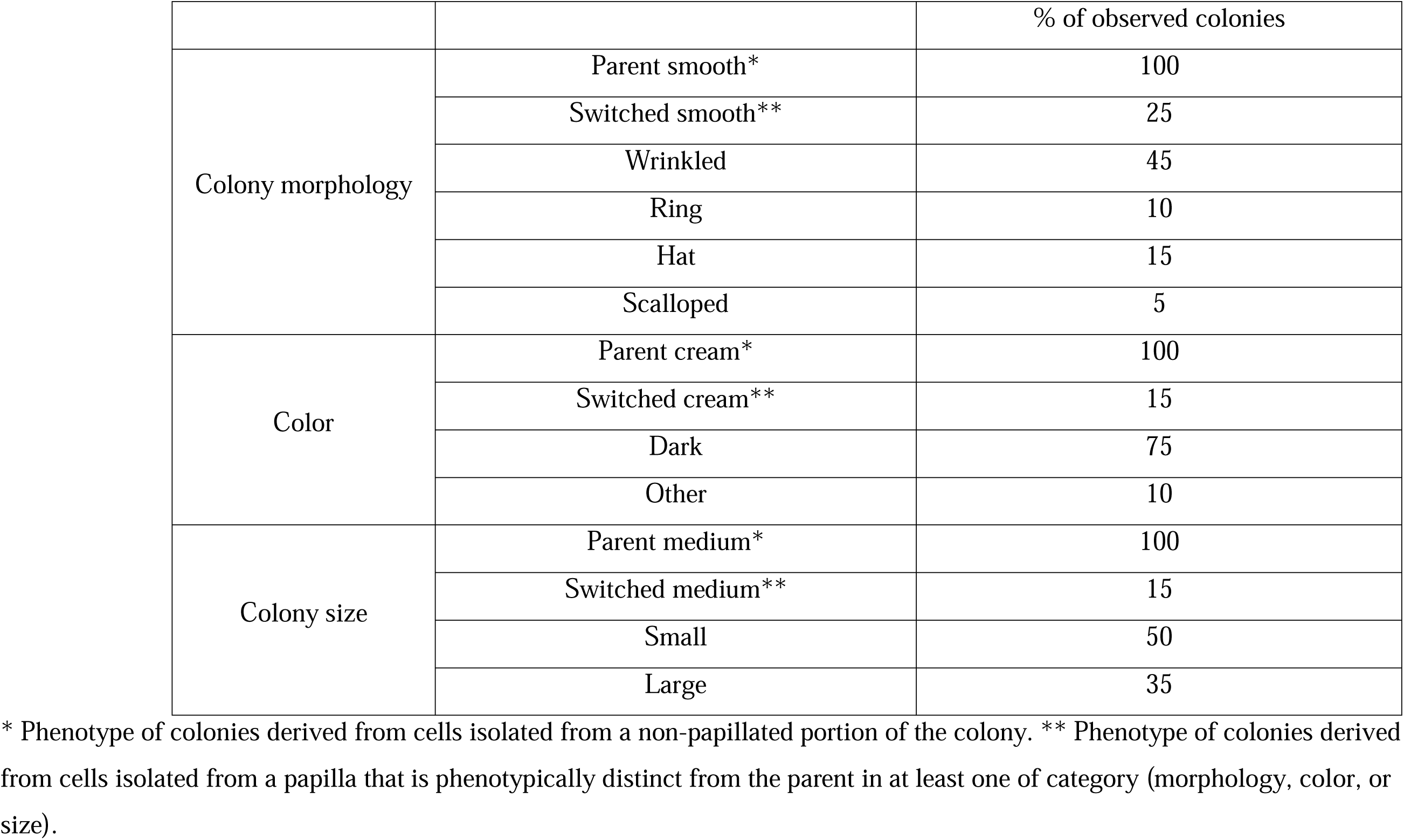
Colony Morphology Differences.

**Table 3:** Phenotypic Progression After Initiation.

| Initial Phenotype | Subsequent Phenotype | % of observed colonies |
| --- | --- | --- |
| wrinkled | wrinkled | 98.8 |
|  | smooth white | 0.91 |
|  | smooth opaque | 0.29 |
| ring | ring | 98.6 |
|  | smooth white | 0.85 |
|  | smooth opaque | 0.44 |
|  | wrinkled | 0.07 |
| smooth opaque | smooth opaque | 99.1 |
|  | ring | 0.80 |
|  | wrinkled | 0.15 |

We noted that with longer incubations, such as shown in Figure 1, multiple papillae form on the same colony and we have observed >5 individual papillae on the same colony. Purification of cells from different papillae derived from the same colony could give rise to different phenotypic variants, suggesting that papillae are arising independently of each other (Figure 1C). These results demonstrate that papillae formation is associated with phenotypic changes not observed in aged matched non-papillated regions. Furthermore, we have examined numerous additional homozygous mutants, including members of the *RIM101* pathway and mutants from a Tn7-insertion library [35] and have not observed papillae formation suggesting that papillae formation is not a general phenomenon. Thus, loss of Mds3 leads to papillae formation in a time-dependent yet apparently stochastic manner.

### Papillae formation linked to CMPS

Colonies from *mds3*Δ*/*Δ*-*derived papillae are phenotypically similar to the previously reported clinical isolates that have undergone colony morphology phenotypic switching (CMPS) [18]. Although CMPS has been associated with the transition from the commensal state to a virulent state, the regulation of CMPS has not been well studied.

Thus, we wanted to determine if the *mds3*Δ*/*Δ-papillae associated switching was in fact a CMPS change as the *mds3*Δ*/*Δ mutant gives us a genetic handle to elucidate the mechanism(s) governing CMPS. In addition to penetrance, described above, two key characteristics define CMPS: continual low frequency switching and additional phenotypic changes unlinked to the colony phenotype [11,18]. We tested these two criteria to determine if our *mds3*Δ*/*Δ derived papillae represent CMPS.

First, we measured the switching frequency in three independent *mds3*Δ*/*Δ-derived variants each with a different colony morphology: wrinkled, ring, and smooth opaque. Each strain was grown overnight and plated on YPD at a concentration of ∼200 colonies/plate and incubated at 37°C for 2 days. In all three cases new morphological variants, based on colony morphology, size, and/or color, were observed with a frequency of ∼10^-2^ (Table 2). For each strain at least two distinct variants arose, and two strains yielded an *mds3*Δ*/*Δ parental colony morphology. These results demonstrate that once switching is initiated, via papillation, switching continues at low level and that switching may be reversible back to the original phenotype. While our switching frequency is ∼10x higher than that observed previously [18], these results support the idea that Mds3 inhibits CMPS.

Second, we tested whether fluconazole sensitivity varied independently to colony morphology. As the wrinkled colony morphology was the most frequent colony phenotype observed, we compared a collection of nine independent *mds3*Δ*/*Δ variants with the wrinkled morphology for fluconazole sensitivity by disk diffusion assay and quantified sensitivity and tolerance using diskImageR (Figure 2 and Table 4) [34]. The inhibitory concentration of fluconazole, measured by RAD_20_, did not vary markedly among the wild-type, the original *mds3*Δ*/*Δ mutant, and nine *mds3*Δ*/*Δ-derived wrinkled isolates (Table 4). Wild-type and the *mds3*Δ*/*Δ mutant gave a RAD_20_ of 13 mm and 14 mm respectively. However, a fluconazole evolved resistant strain (AMS4058) from the same genetic background had a RAD_20_ of 5 mm. The wrinkled *mds3*Δ*/*Δ variants gave a RAD_20_ range from 11-16 mm. Thus, neither loss of Mds3 nor the wrinkled colony phenotype in the *mds3*Δ*/*Δ background dramatically impacted on fluconazole sensitivity. However, there was marked variation in tolerance as measured by FoG_50_. Wild-type and the *mds3*Δ*/*Δ mutant had a FoG_50_ of 0.08 and 0.14 respectively and AMS4058 was completely tolerant. This indicates that loss of Mds3 results in slightly more tolerance to fluconazole despite having a similar sensitivity to the drug compared to wild-type cells. However, the wrinkled *mds3*Δ*/*Δ variants showed marked variation in FoG_50_ ranging from 0.07 (similar to wild-type) to 0.68. In total, the low level of continued switching and tolerance variability support the idea that papillae arising on the surface of *mds3*Δ*/*Δ mutant colonies represent CMPS and that Mds3 is a negative regulator of CMPS in *C. albicans*. To our knowledge Mds3 represents the first established regulator of CMPS.

**Table 4:**
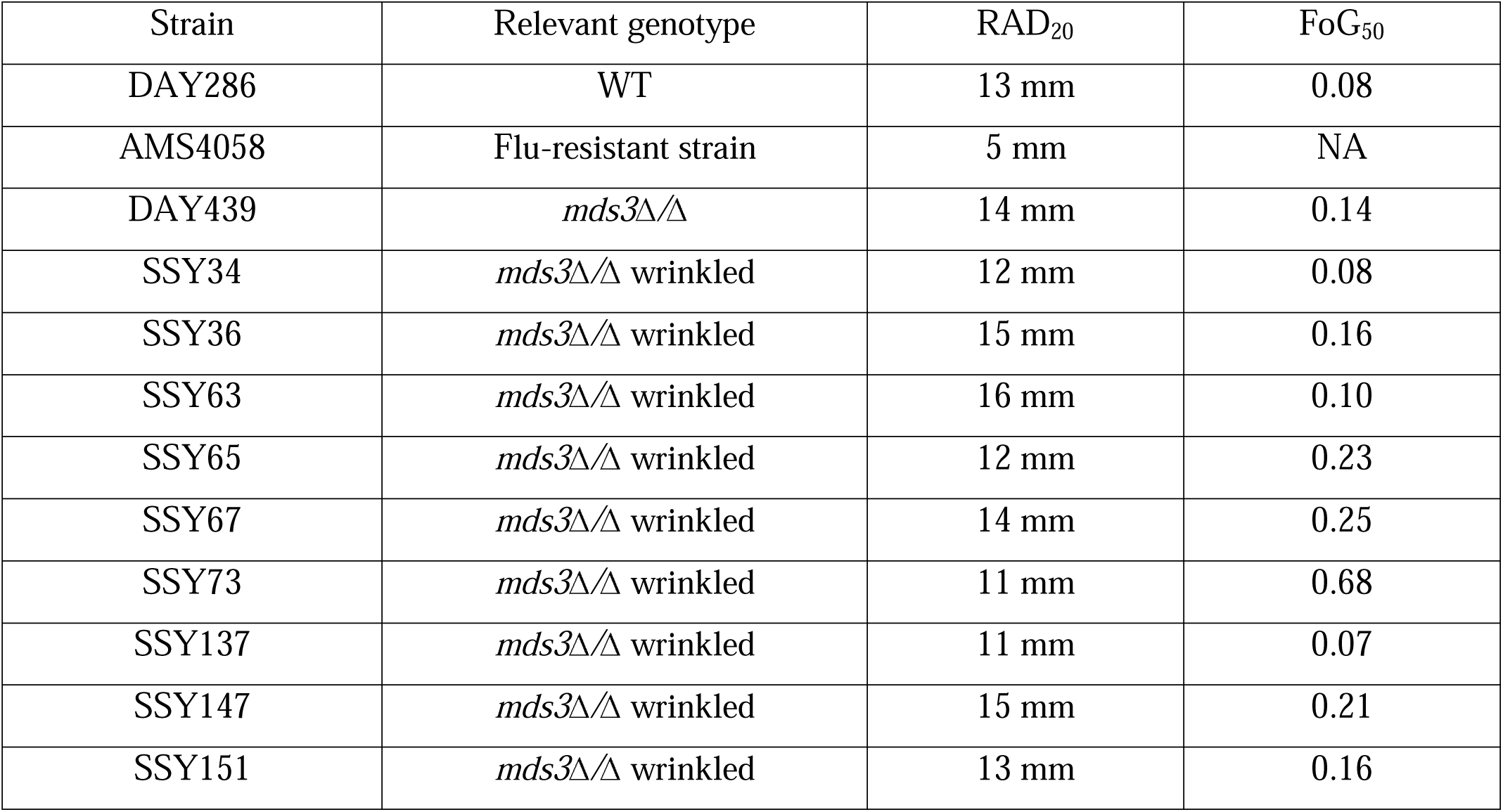
Fluconazole Sensitivity and Tolerance.

### CMPS Is Due to Alterations in the TOR Pathway

We demonstrated previously that Mds3 functions in the TOR growth control pathway in *C. albicans* and *S. cerevisiae* [36]. Thus, we wanted to determine if papillae formation and CMPS is due to Mds3 TOR-dependent function(s) or to a distinct role for Mds3. Loss of Mds3 leads to rapamycin resistance in *C. albicans* [36]. If the CMPS observed in *mds3*Δ*/*Δ mutants reflect a change in the TOR pathway, we reasoned that *mds3*Δ*/*Δ switched isolates derived from the papillae may have altered rapamycin sensitivities compared to *mds3*Δ*/*Δ age matched non-switched cells derived from a non-papillated part of the colony (Figure 3). Of the six *mds3*Δ*/*Δ-derived CMPS:non-switched pairs tested, all of the age matched non-switched isolates behaved like the parental *mds3*Δ*/*Δ strain and were resistant to rapamycin. However, the switched isolates were more sensitive to rapamycin than their presumably clonal non-switched partners (Figure 3). Five of the switched isolates were at least 100x more sensitive to rapamycin with two isolates, SSY34 and SSY137, showing rapamycin sensitivity similar to wild-type strains.

**Figure 3:**
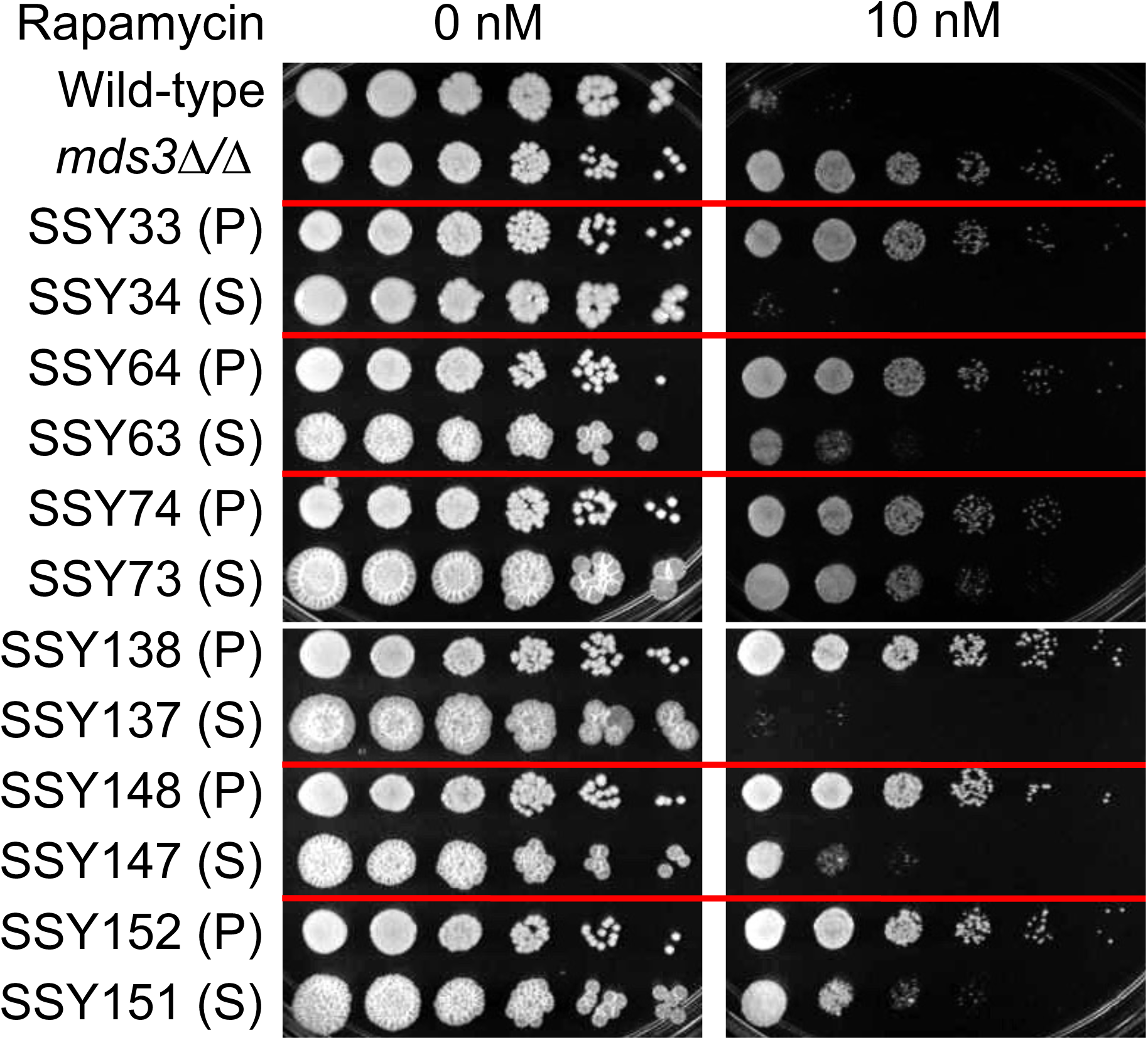
Rapamycin sensitivity in *mds3*Δ*/*Δ switched strains. Growth of wild-type (DAY185), *mds3*Δ*/*Δ (DAY1118), and *mds3*Δ*/*Δ non-switched (P) and switched (S) isolates derived from DAY417 and DAY439 on spider medium containing 0 nM (solvent only) or 10 nM rapamycin.

Previously, we found that the rapamycin sensitivity of the *mds3*Δ*/*Δ mutant was rescued by the rapamycin resistant *TOR1-1* allele [36]. To see if *TOR1-1* affects CMPS, we measured papilli formation of *TOR1-1/tor1*Δ and *TOR1/tor1*Δ strains with and without *MDS3* (Figure 4). Following fourteen days of incubation, wild-type colonies did not develop papilli. However, ∼5% of *TOR1-1/tor1*Δ colonies did give rise to papilli over this time frame. Because the *TOR1-1* strain contains a single copy of *TOR*, we analyzed the *TOR1/tor1*Δ heterozygous mutant to determine if the low level papilli formation observed in the *TOR1-1/tor1*Δ background was an issue of gene dosage or an attribute of the *TOR1-1* allele. Similar to wild-type, the *TOR1/tor1*Δ strain did not show papilli on colonies. These results suggest *TOR1-1* may have a minor effect on CMPS. In the *mds3*Δ*/*Δ background, the *TOR1/TOR1* and *TOR1/tor1*Δ heterozygous strains behaved similarly with ∼30% of colonies having papilli (Figure 4). However, in the *mds3*Δ*/*Δ *TOR1-1/tor1*Δ strain, a marked increase in papilli formation was observed with >90% of colonies having papilli. These results demonstrate that *TOR1-1* exacerbates papilli formation observed in the *mds3*Δ*/*Δ background.

**Figure 4:**
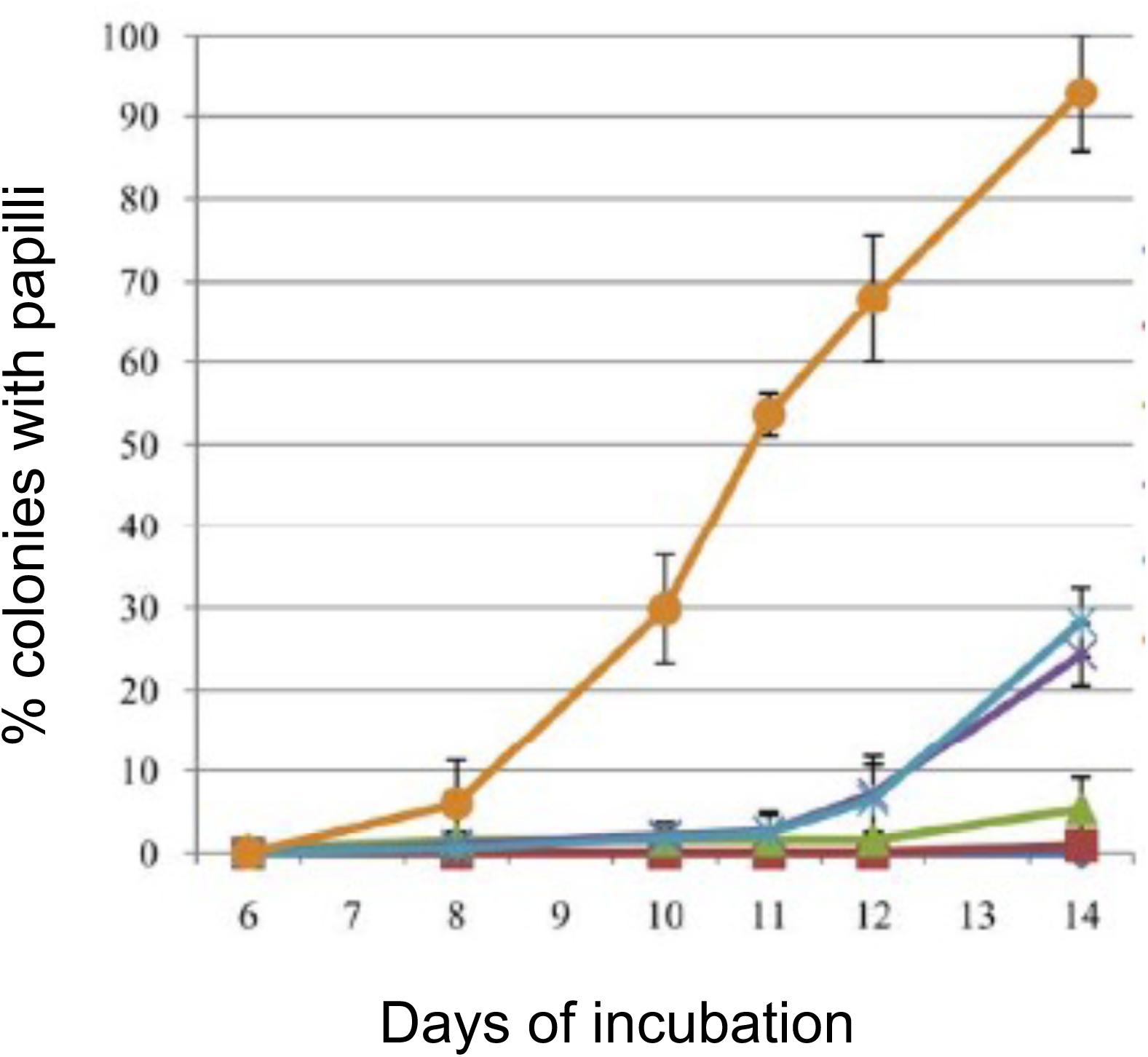
Effect of the *TOR1-1* allele on *mds3*Δ*/*Δ-dependent papilli formation. Overnight cultures from the wild-type (DAY185, blue diamonds), TOR1/Δ (DAY1123, red squares), TOR1-1/Δ (DAY1125, green triangles), mds3Δ/Δ (DAY1118, purple x), mds3Δ/Δ TOR1/Δ (DAY1122, light blue asterisks), and mds3Δ/Δ TOR1-1/Δ (DAY1124, orange circles) strains were diluted in PBS and plated on YPD. Plates were incubated 48 hours at 30°C and then at room temperature for two weeks. The results are representative of two independent experiments and the standard error is shown.

Finally, we analyzed several additional mutations affecting TOR pathway members for papillae formation. Sch9 is activated by TOR kinase and promotes growth whereas Sit4 is activated in the absence of TOR signaling and promotes starvation responses [36–39]. We observed that colonies derived from the *sch9*Δ*/*Δ mutants, but not the *sit4*Δ*/*Δ mutant, generated papillae with prolonged incubation (Figure 5). Purification of cells from *sch9*Δ*/*Δ-derived papillae gave rise to phenotypically switched colonies analogous to those observed from Mds3 deficient strains (data not shown). Based on these pharmacological and mutant studies, we conclude that the TOR pathway plays a key role in CMPS in *C. albicans*.

**Figure 5:**
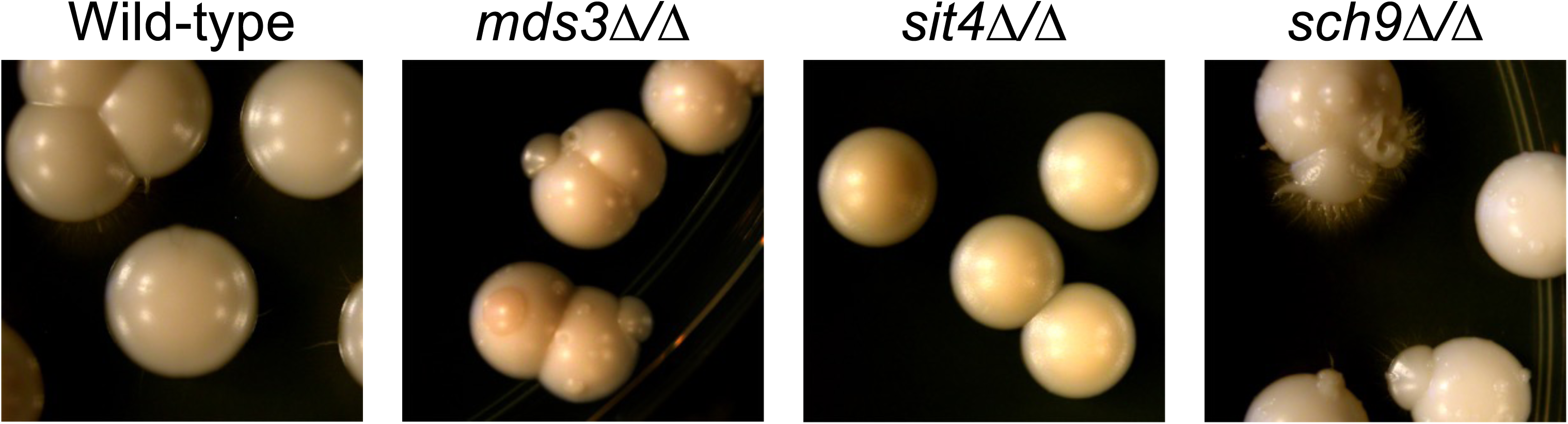
Papilli formation on colonies of TOR pathway mutants. DAY185 (wild-type), *mds3*Δ*/*Δ (DAY1118), *sit4*Δ*/*Δ (DAY971), and *sch9*Δ*/*Δ (DAY1225) were incubated 2 days at 30°C and then stored another 10 days on the bench to allow papilli formation. Pictures are a representative region of a single plate per strain.

### CMPS and TOR in Wild-type and Clinical Isolates

Our studies implicate the TOR pathway as a key regulator of a CMPS in *C. albicans*. However, this work depended on mutant backgrounds which may not reflect how CMPS occurs clinically. To test a potential link between the TOR pathway and clinical CMPS, we needed to establish *in vitro* conditions to promote CMPS. Previously, CMPS was observed in the laboratory using a single strain background and was induced by UV irradiation [18,40], which may not reflect CMPS in a clinical setting. Thus, we obtained a collection of commensal *C. albicans* isolates [31,41] and tested whether they undergo CMPS and, if so, if this CMPS related to the TOR pathway.

Neither the parental wild-type lab strain nor clinical isolates formed papillae on solid medium with prolonged incubation (Figures 1A and data not shown). Since CMPS was only observed clinically concomitant with increasing *C. albicans* population size, we moved to a liquid culture assay in which we could more readily alter population sizes, nutrient concentrations, etc. After overnight growth in YPD at 37°C our wild-type lab strain did not yield colonies reminiscent of CMPS (Figure 6A, left). We did note a few colonies with slight ‘wrinkly’ phenotype (Figure 6A left arrows), but purification of cells from these colonies did not recapitulate that phenotype. This suggests that these cells were inducing some filamentous growth due to incubation at 37°C and not CMPS. In contrast, after two weeks of incubation at 37°C, CMPS variants were readily identified with both parental-like colonies and phenotypic variants occurring (Figure 6A, right).

**Figure 6:**
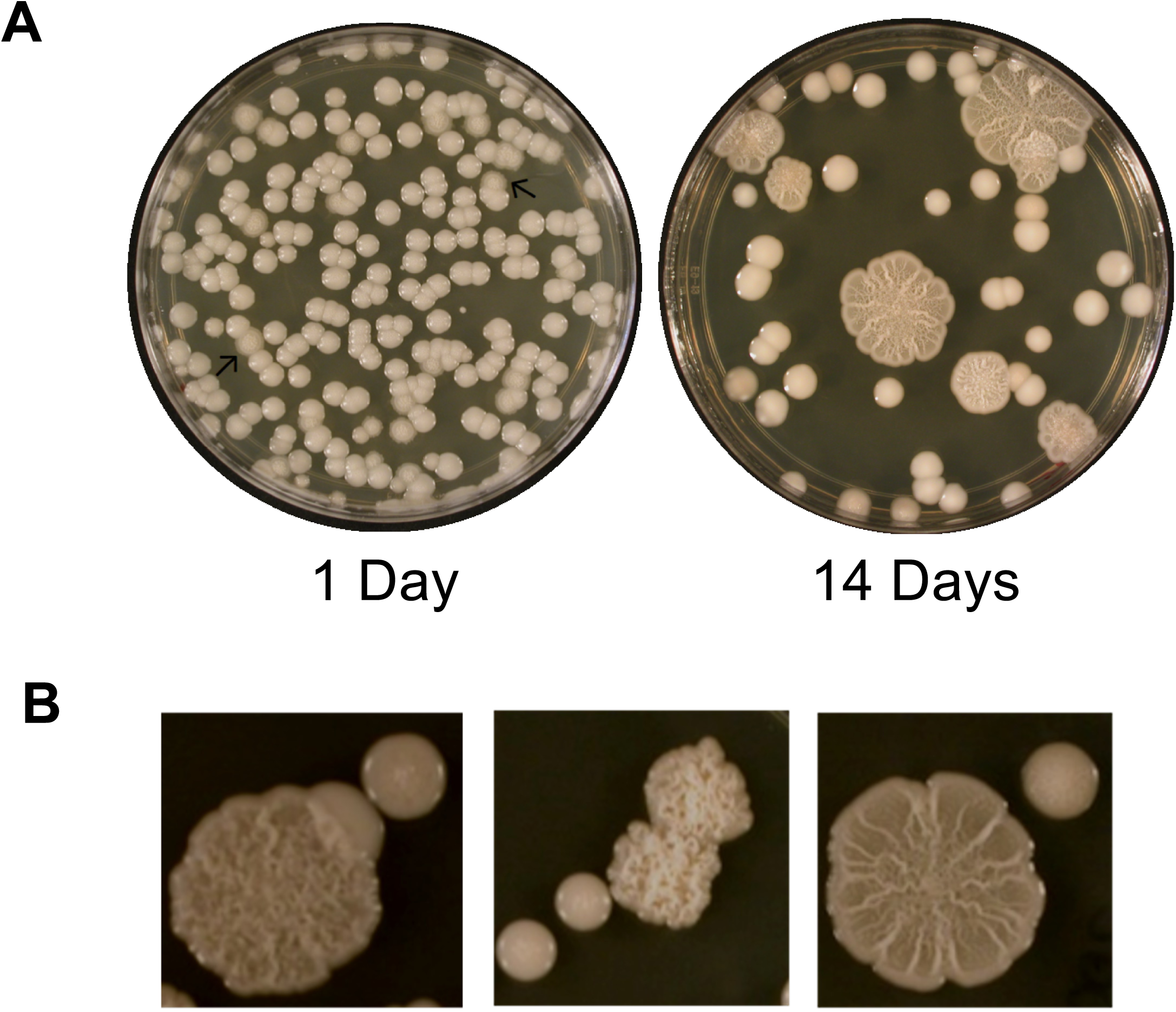
Switching in wild-type *C. albicans* strains. A. The lab wild-type strain (DAY185) was plated onto rich solid medium after overnight and fourteen days incubation at 37°C in liquid rich medium. B. Three independent clinical isolates plated onto rich solid medium after fourteen days incubation at 37°C in liquid rich medium. Photographs were taken after two days growth at 37°C.

Cells restreaked from these phenotypically switched colonies gave rise to colonies with the same phenotypic morphology demonstrating that the phenotypic switch was penetrant. We also observed CMPS in all clinical isolates tested following prolonged incubation in liquid medium at 37°C (Figure 6B). Importantly, we observed both parental and switched colonies in each case. In all cases, we noted a dramatic loss in plating efficiency with increased time of incubation and observed that onset of CMPS was concomitant with this decrease in viability.

Because our liquid cultures were inoculated with a single colony, the switched and non-switched colonies obtained after plating must be extremely close genetic relatives and likely clones. Thus, we could test whether CMPS in these backgrounds reflect changes in the TOR pathway by comparing switched and non-switched pairs. We predicted that switched colonies would show differences in rapamycin sensitivity compared to non-switched sister colonies. To test this hypothesis, we screened switched and non-switched colonies obtained from the same plate for rapamycin sensitivity (Figure 7). We found that rapamycin sensitivity varied markedly in the non-switched isolates from extremely sensitive (DAY979P) to resistant (DAY965P). This, not surprisingly, indicates that different genetic backgrounds have different sensitivities to rapamycin.

**Figure 7:**
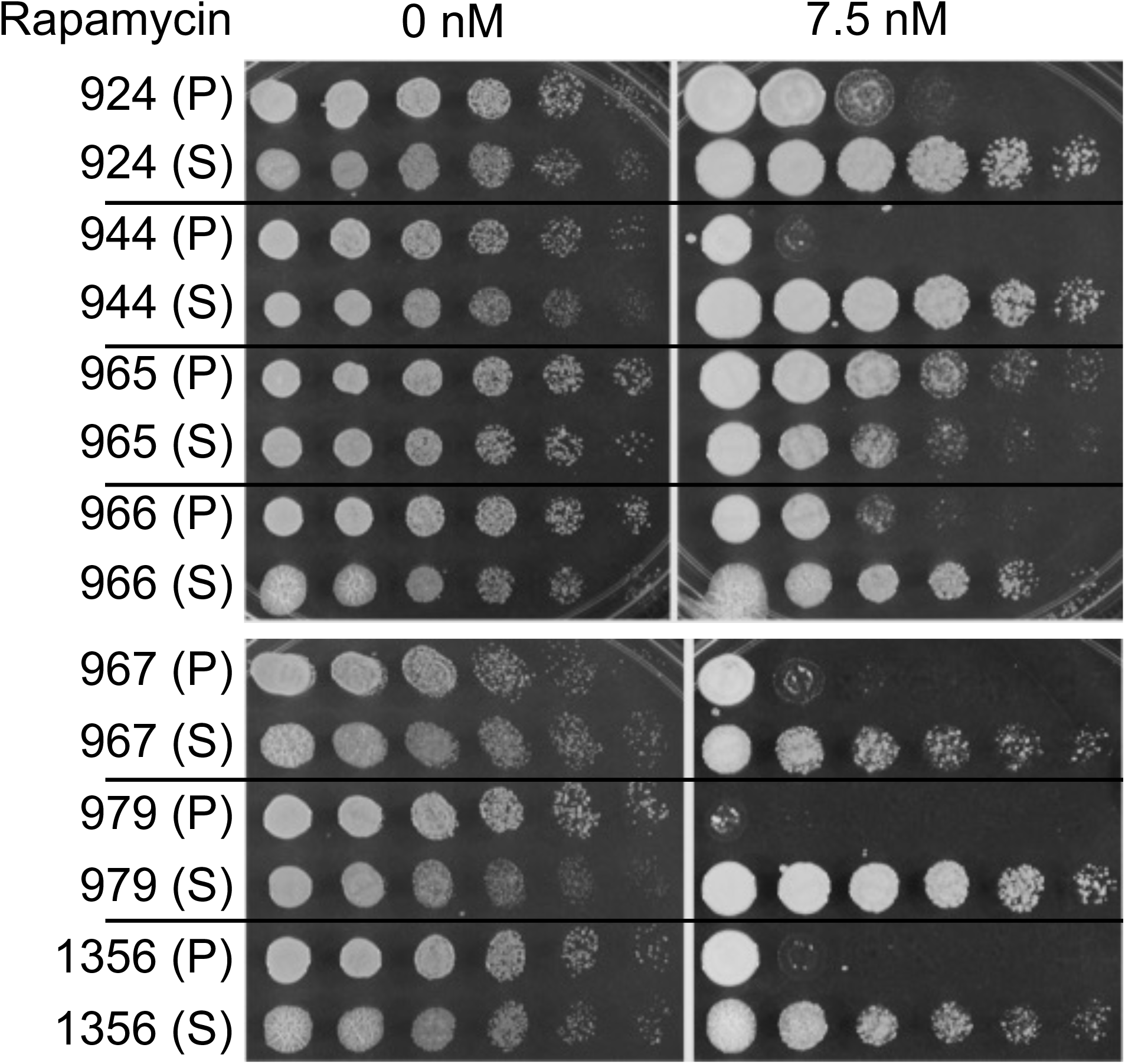
Rapamycin sensitivity in wild-type switched strains. Growth of wild-type (non-switched (P) and switched (S) isolates derived from DAY924, DAY944, DAY965, DAY966, DAY967, DAY979, and DAY1356 on spider medium containing 0nM solvent only) or 7.5nM rapamycin.

However, of the seven switched:non-switched pairs tested, six of the switched isolates were at least 100x more resistant to rapamycin compared to their non-switched partner. The remaining switched isolate (DAY965S) appeared to be slightly more sensitive than its matched sister and is from the most resistant background. Overall, these results support the conclusion that the TOR pathway is a key regulator for CMPS in *C. albicans*.

## Discussion

The ability to generate and maintain phenotypic diversity is a critical aspect of a species’ ability to survive in an ever-changing environment. This is also true for microbes that colonize and infect mammalian hosts, such as *C. albicans* and *H. pylori*. In most eukaryotes, genetic diversity, which is generated by sexual reproduction, is a primary source of phenotypic diversity. Although *C. albicans* can mate and is able to undergo a parasexual cycle, these events appear to be rare in nature and do not appear to be sufficient to generate or maintain genetic diversity [42]. However, the process of colony morphology phenotypic switching (CMPS) generates colonies with distinct yet penetrant phenotypes. In addition to yielding an altered colony morphology, CMPS variants show differences in antifungal drug resistance, protease secretion, and other phenotypes, which are not directly related to colony phenotype [11,19,43]. CMPS occurs *in vivo* and CMPS variants have been isolated from both oral and vaginal cavities [13] [18]. Thus, CMPS functions to generate phenotypic diversity in this asexual eukaryote and we demonstrated that CMPS is associated with alterations of the TOR pathway using both genetic and pharmacological approaches.

We found that CMPS requires prolonged incubation to be observed in both wild-type and mutant backgrounds. We cannot rule out the possibility that CMPS occurs constitutively, but under rapid growth conditions the wild-type phenotype outcompetes switched strains. Indeed, a previous study by Soll et al found that rare CMPS isolates could be found within normal colonies of the 3153A background following a seven-day incubation at room temperature [18]. Thus, starvation may induce CMPS or allow CMPS isolates to outcompete non-switched sisters. These prolonged growth conditions are expected to promote stationary phase or dormancy, however our results indicate that at least low levels of growth occurs. This idea is best observed on solid medium in the *mds3*Δ*/*Δ and *sch9*Δ*/*Δ mutants, which give rise to actively growing papillae with prolonged time. We predict that there is a change within a specific cell or sub-set of cells that promote growth, presumably feeding on dead sister cells, within the colony.

Importantly, distinct papillae from the same colony behave differently suggesting that independent events are occurring within the population driving evolution. In liquid culture, we observed a profound loss in viability concomitant with onset of CMPS; these dead cells could provide the nutrients allowing growth of CMPS variants. Conversely, CMPS may generate a sub-set of cells that are resistant to this death but are not actively growing.

Previously, we found that Mds3 and Sit4 physically interact and showed that Mds3 functions in the starvation arm of the TOR pathway [36]. The fact that the *mds3*Δ*/*Δ and *sch9*Δ*/*Δ mutants, but not the *sit4*Δ*/*Δ mutant, behave similarly in CMPS indicate that Mds3 functions more generally in the TOR pathway. Indeed, our unpublished transcriptional analyses support the idea that Mds3 acts more generally in both the growth and starvation arms of the TOR pathway.

How does the TOR pathway promote CMPS? Initiation of cellular growth is controlled by signal transduction systems which sense and respond to nutrient availability. The TOR kinase, an essential protein conserved throughout the eukarya, is regulated in response to nitrogen availability. When sufficient nitrogen is available, TOR kinase is activated and phosphorylates targets that stimulate growth, including Sch9, the ortholog of mammalian S6-Kinase I [44,45]. TOR also phosphorylates Tap42 which inhibits Sit4, a serine-threonine phosphatase required for starvation responses like autophagy and nitrogen-catabolite repression [37]. When nitrogen is limited, TOR kinase is inactive leading to reduced Sch9 activity and dephosphorylation of Tap42, promoting Sit4-dependent starvation responses. As noted above, in liquid culture, plating efficiency drops precipitously. These dead cells may provide sufficient nitrogen that promote a low-level of growth and as these cells divide mutations accumulate some of which alter TOR activity in these reduced nitrogen conditions, altering rapamycin sensitivity, and resulting in CMPS. Another possibility is based on the role of Sch9 in chromosome stability. In *C. albicans*, Sch9 associates with the centromere and loss of Sch9 is associated with increased chromosome loss [39]. Previous studies of clinical CMPS isolates showed that aneuploidies were common [46,47]. This latter hypothesis provides clear predictions to explain the phenotypic diversity (differences in specific aneuploidies), continued low level switching (changes in aneuploidies), and reversion to the wild-type phenotype (restoration of the diploid genomic content).

One confounding issue is that switched strains derived from *mds3*Δ*/*Δ colonies were more sensitive to the TOR inhibitor rapamycin, whereas switched strains derived from wild-type strains were more resistant to rapamycin. We predict that this difference is due to the already altered TOR pathway in the *mds3*Δ*/*Δ mutant background. Loss of *mds3*Δ*/*Δ causes increased rapamycin resistance [36], which may be disrupted when the TOR pathway changes promoting CMPS occur. Conversely, in wild-type backgrounds, those same changes promote resistance.

CMPS promotes phenotypic variation in the asexual fungus *C. albicans* and has been observed clinically for almost 100 years. Here we provide evidence that CMPS is controlled via the TOR growth control pathway which we believe is the first clear regulatory system described for this phenomenon. In conjunction with previous studies, our results suggest that CMPS is not a stochastic process, although the outcome of CMPS may be stochastic. We also note that TOR is conserved throughout eukaryotic evolution and TOR dysfunction is strongly implicated in a number of cancers that also readily generate phenotypic variants [27], suggesting our work may be applicable beyond fungal pathogenesis.

## Acknowledgments

This work was supported by the Investigators in Pathogenesis of Infectious Disease Award from the Burroughs Wellcome Fund to D.A.D. We are indebted to Dr. Judith Berman, Dr. Anna Selmecki, current and former members of the Davis lab, as well as colleagues at the University of Minnesota for helpful feedback and guidance over the course of these studies. We are also indebted to Dr. Selmecki for sharing AMS4058 with us. We are indebted to the Tissue Procurement Facility, Fairview University Medical Center, Minneapolis, and Matthew Larson in the Department of Medicine who were collectively responsible for the primary interactions with the patients. We greatly appreciate the willingness of many anonymous patient donors who have provided samples for research purposes.

